# Telomeric dysfunction triggers an unstable growth arrest leading to irreparable genomic lesions and entry into cellular senescence

**DOI:** 10.1101/442533

**Authors:** Sabrina Ghadaouia, Marc Alexandre Olivier, Aurélie Martinez, Nicolas Malaquin, Guillaume B Cardin, Francis Rodier

## Abstract

Replicative senescence is the permanent growth arrest caused by gradual telomere attrition occurring at each round of genome replication. Critically shortened telomeres lose their protective shelterin complex and t-loop structure revealing uncapped chromosome ends that are recognized as DNA double-strand breaks causing a p53-dependent DNA damage response (DDR) towards proliferation arrest. Because telomeres are heterogeneous in length within a single cell, the number of short telomeres necessary for senescence onset remains ill defined. Using controlled Tin2-mediated shelterin inactivation, we show that telomere uncapping is not sufficient to trigger senescence. While uncapping generates expected telomeric DNA damage detection, the associated weak DDR allows a rapid bypass of the primary growth arrest and re-entry into the cell cycle despite dysfunctional telomeres. During the ensuing mitosis, fused telomeres lead to additional DNA breaks and to genomic instability including chromosomes bridges or micronuclei, which sustain a secondary entry into stable growth arrest. The loss of p53 prevented both primary and secondary growth arrest, leading to amplified genomic instablility. Our results support a new multistep model for entry into telomere-mediated replicative senescence in normal cells, which is not directly induced by telomere uncapping, but rather by an amplification of DNA lesions caused by telomere fusions that leads to permanent irreparable genome damage.

## INTRODUCTION

Replicative cell senescence was described based on the observation that cultured cells have a limited proliferation potential leading to an irreversible growth arrest associated with telomere shortening ^1,2^. Cellular senescence is involved in several physiological mechanisms such as aging, wound healing, cancer suppression, and cancer promotion ^3^.

Telomeres are non-coding repetitive DNA sequences at the extremity of every chromosome and play an essential role to protect the free DNA ends from exonuclease-mediated degradation or end-to-end fusions ^4^. Telomeric end protection is provided by the telosome, or shelterin complex, formed by 6 subunits (TRF1, TRF2, Tin2, Rap1, POT1, and TPP1) that assemble on the telomeric DNA repeats ^5,6^. The complex forces telomeres into a t-loop structure, where the single-stranded DNA overhang is hidden ^5,6^. Telomere repeat factors (TRF) 1 and 2 bind directly to telomeric double-strand DNA, while protection of telomere 1 (POT1) binds simultaneously to the single-strand DNA overhangs and to TRF2. Repressor / activator protein 1 (Rap 1) interacts with TRF2, while TINT1-PTOP-PIP1 (TPP1) binds to POT1. Finally, TRF1-2 interacting nuclear protein 2 (Tin2) creates a bond between TRF1-TRF2 and the POT1-TPP1 complexes, thereby bridging shelterin subunits that are attached to double-stranded DNA and those that are attached to single-stranded DNA. Altogether, these proteins ensure the stability of the t-loop, and therefore contribute to the protection of the chromosome extremity. Additionally, the shelterin complex can provide active inhibition of the DNA damage response (DDR), which is responsible for cell cycle checkpoint signalling and for DNA repair at telomeres, thus preventing untimely activation of cell cycle checkpoints or chromosome end-to-end fusions during DNA replication ^4,7^. For example, TRF2 ^4^ and POT1-TPP1 dimer ^7^ prevent the activation of ATM and ATR, the two main upstream kinases for DDR, following double-strand breaks (DSB) or single-strand breaks (SSB) respectively. Similarly, direct inhibition of DNA repair pathways such as Ku70/Ku80-associated non-homologous end joining (NHEJ), is mediated by TRF2 ^8^ and Rap1 ^9^. Overall, telomeric end protection via the shelterin complex is crucial to ensure genome stability during cell division.

In addition to their role in preventing genome instability, telomeres act as a reflection of cellular aging, as the ‘’end replication problem’’ causes their shortening at every cell division ^10^. Following several rounds of erosion, telomeres become too short to form a proper t-loop and can no longer protect the DNA at chromosome ends ^5^, resulting in a DDR, associated with the formation of telomeric DNA damage foci (DDF) that co-localize with 53BP1 and γH2AX ^11^. This leads to the activation of DDR and p53-mediated cell cycle checkpoints that definitively arrest the proliferation of normal cells, establishing replicative senescence ^11-15^.

Despite this mechanistic understanding of telomere-mediated senescence, a key question regarding the initiation and maintenance of a stable replicative senescence growth arrest in mammalian cells remains unanswered: how many uncapped telomeres are necessary to trigger replicative senescence? This question emerges from the fact that telomeres are heterogeneous in length within single cells ^16^, and the rate of shortening varies significantly between telomeres due to random telomeric DNA breaks ^17^ and to the end replication problem itself ^10,18^. Overall, the rate varies between 50 and 150 bp per replication, and the average telomere length prior to senescence is 1–2 kb ^19^. Therefore, it remains impossible to predict which telomere will be shortened in a cell nearing replicative senescence.

Surprisingly, mammalian telomeres shorter than 1 kb are observed in proliferating cells ^19^. These normal cells were also able to proliferate with telomeres that were too short to be recognized by probe hybridization ^21^, suggesting that mammalian cells can possibly tolerate a certain amount of dysfunctional telomeres. Similarly, stable short telomeres were observed in proliferating fibroblasts, suggesting that a limited number of short telomeres did not prevent proliferation ^21,22^. It has been suggested that approximately 5–10 dysfunctional telomeres, perhaps even more, are required to trigger replicative senescence ^21,22^.

Taken together, it appears that a single short telomere is not sufficient to trigger replicative senescence in mammals, and that cells are able to proliferate while harbouring an as yet undefined number of dysfunctional telomeres. To test cell tolerance for telomere DNA damage, we used a different approach and took advantage of the fact that disruption of telomeric chromatin can be controlled via the inactivation of essential components of the shelterin complex, leading to immediate telomere uncapping and DDR activation ^4,23,24^. We used a well characterized model of normal human fibroblasts harbouring an inducible dominant negative mutant of Tin2 that cannot bind TRF1, Tin2DN (Tin2-15C in ^25,26^). We tested to what extent cells could tolerate direct telomere uncapping, irrespective of telomere length, and found that normal human cells were tolerant of widespread telomere uncapping. While t-loop disruption triggered a telomeric DDR that eventually lead to cellular senescence, a time course follow-up of entry into senescence revealed a multistep relationship with telomeric damage and growth arrest. We reveal a cell cycle checkpoint bypass in the presence of telomeric DDF that causes secondary breaks and genomic instabilities, and demonstrate that a permanent senescence growth arrest in normal cells is associated with irreversible genome damage caused by a telomeric fusion-bridge-break cycles.

## MATERIAL AND METHODS

### Cell culture

Normal human fibroblasts (HCA2 and BJ-U) were cultured at 37°C with 20% O_2_ and 5% CO_2_ in DMEM media (Multicell, Wisent Inc.) with 8% FBS (Fetal Bovine Serum) and 5% antibiotics: penicillin and streptomycin (Wisent Inc.). IMR90 cells, another type of normal human fibroblasts, were cultured in the same media, but at 5% O_2_. Cells were then treated with (Dox ON) or without (Dox OFF) Doxycycline at 30 ng/mL for 168 h with fixation at every 24 h for analysis. For our low serum studies, cells were cultured with 0.1% FBS DMEM media.

### Live Imaging

We performed live-imaging acquisition using IncuCyte ZOOM system, a microscope integrated to the incubator to obtain real time cell imaging. We used BJ-U TR4s Tin2DNs +/- shp53 H2B GFP cells to visualize nuclei and follow our cells throughout induction. We took pictures every hour for 168h in order to observe mitosis.

### Genetic modifications of primary cells

We used two types of primary normal human fibroblasts: HCA2 and BJ-U. Using Invitrogen’s Gateway technology (Campeau et al., 2009), we infected them with lentivirus containing a Tetracyclin Repressor (TetR), creating HCA2 TR3s and BJ-U TR4s primary cells. Still using the Gateway system, we infected HCA2 cells with a lentivirus containing p16 to obtain HCA2 TR3s p16s, primary normal human fibroblasts with the ability to express p16 when induced with Doxycycline. Similarly, we created BJ-U TR4s Tin2DNs and BJ-U TR4s shTin2s. In addition, BJ-U TR4s Tin2DNs cells were infected with constitutively expressed lentivirus (Doxycycline independent) to create the following cells: BJ-U TR4s Tin2DNs H2B-GFP, BJ-U TR4s Tin2DNs shp53 and its control BJ-U TR4s Tin2DNs shGFP.

### Immunofluorescence

Cells were cultured in 4 well chamber slides (Nunc, Penfiled) then fixed in 10% formalin (Sigma). Cells were then permeabilised with PBS containing 0.25% Triton X-100 and fixed in a blocking buffer composed of 1% BSA (Jackson Immuno Research) and 4% donkey serum (Jackson ImmunoResearch) in PBS. Primary antibodies were incubated at 4°C overnight in the blocking buffer, then washed for 30 min at room temperature. They were used to detect 53BP1 (NB100-304, Novus Biologicals), diluted at 1/2000; gH2AX S139 (05-636-I, Millipore) at 1/2000; and p16 (JC8, Santa Cruz) at 1/300. Secondary antibodies (Alexa Fluor 488nm or 647nm) were incubated for 1 h at room temperature, and diluted 1/800 in blocking buffer. Finally, slides were washed and mounted in ProLong Gold Antifade reagent with DAPi (Life Technologies). Images were acquired with a Carl Zeiss Axio Observer Z1 fluorescence microscope, and quantification was performed with Axio Vision software. First, DAPI staining allowed us to defined nuclei as primary objects. We then identify 53BP1 staining that is within the nuclei (i.e., primary object) and with an intensity above a defined intensity threshold and in between a size range. Finally, we obtain the number of 53BP1 foci in each nuclei.

### Western Blot

Cells were lysed in MPER reagent (Thermo Scientific) with a cocktail of protease and phosphatase inhibitors (Sigma) on ice, then centrifuged at 4°C for 15 min and sonicated at 4°C for 10 min. Protein samples were quantified using the BCA protocol (Thermo Scientific) and mixed with BlueJuice gel loading buffer (Invitrogen) and DTT (Sigma) before a 5 min incubation at 95°C. Samples were then loaded and separated in 4-15% precast SDS PAGE gels (BioRad) for1 h at 110 V before transfer onto a PVDF membrane (Immobilon) at 110 V at 4°C for 1 h. The membrane was then blocked in 2% BSA (Sigma) for 1 h at room temperature and incubated overnight at 4°C with the primary antibody diluted in 2% BSA. Mouse antibody against pATM-S1927 (#4526) was diluted 1/2000 and the rabbit antibody against pchck2-T68 (#2661) at 1/1000 were from Cell Signalling. Mouse antibodies against p53 and p21 from BD Pharmingen were diluted 1/200. After washing and incubation with the secondary HRP antibody, proteins were revealed with chemiluminescence (ECL Western blotting substrate, Thermo).

### EdU

Cells were seeded in 4 well chamber slides (Nunc, Penfiled) and pulsed with the modified thymidine analogue EdU for 24 h, then fixed in 10% formalin (Sigma). EdU was then stained using the Click-iT protocol (Invitrogen) with the Alexa Fluor 647 dye. Slides were washed and mounted in ProLong Gold Antifade reagent with DAPi (Life Technologies). Images were acquired with a Carl Zeiss Axio Observer Z1 fluorescence microscope, and quantification was performed with Axio Vision software.

### LiCor

Cells were cultured in 96 well plates and fixed in formalin (Sigma) before staining with Draq5 diluted at 1/10000 in PBS for 1 h. Plates were read with Odyssey Li-Cor plate reader (Mandel).

### FACS

Cells were cultured in 6 well plates, trypsinized, centrifuged, fixed in 70% ethanol then stained with PI (Sigma) in 0.2% TritonX-100 with RNase A for 30 min. They were then sorted according to their DNA content with FACS. Analysis were made with the FlowJow software. We used the algorithm to define DNA content range and therefore separate our cell cycle phases.

## RESULTS

### Telomere dysfunction-induced cell growth arrest is unstable

To study the cell response following telomeric dysfunction, we induced expression of Tin2DN with Doxycycline treatment to open the t-loop and form telomeric DDF *(Figure S1a, b).* Alternatively, we also induced p16 expression to trigger a reference DNA damage-free senescence growth arrest ^28^. The inductions lasted for 168 hours with cells fixed for analysis every 24 hours. We first measured general cell proliferation using a total DNA labelling dye, Draq5, and observed that while p16 expression results in a rapid and stable growth arrest, cells with telomeric damages induced by Tin2DN only slowed their proliferation to a level somewhat in-between control cells (non-induced) and p16-induced cells (*Figure 1a)*.

**Figure 1:**
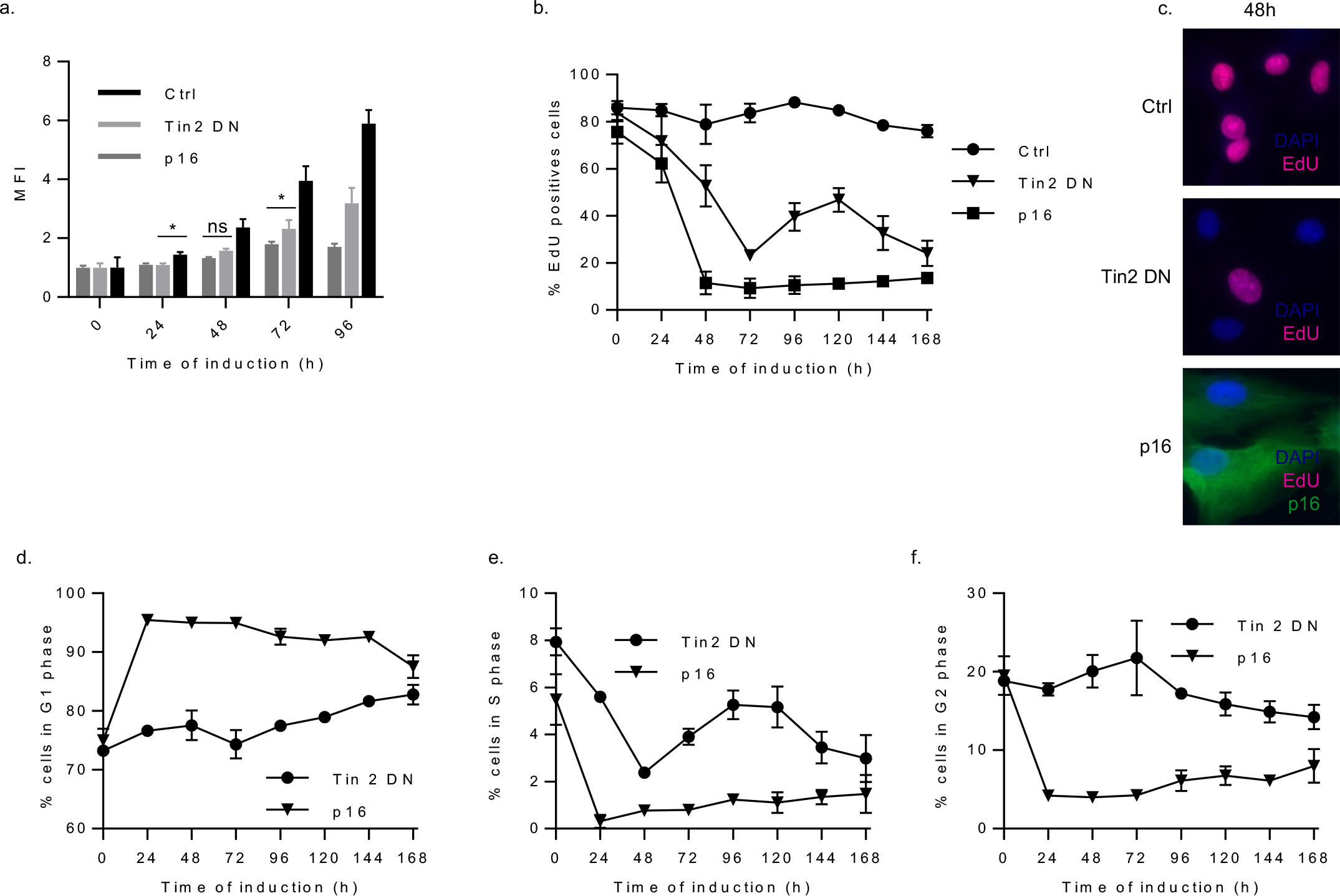
Tin2DN induced growth arrest is unstable. *(a)* Cells were treated with Doxycycline to express p16 or Tin2DN and fixed at different time points. Control cells were not induced. They were stained with Draq5 and the mean fluorescent intensity (MFI) measured with LiCor. Error bars correspond to the standard deviation between the 3 repetitions *(b)* Cells were treated with Doxycycline and EdU was added to the media 24h prior to fixation. Cells were then stained and EdU positive cells quantified. Error bars correspond to the standard deviation between the 5 repetitions *(c)* Representative images of the 48 h time point are shown for each condition. *(d,e,f)* Cells were treated with Doxycycline and trypsinised at different time points. They were then stained with PI and DNA content was quantified with FACS. Here are shown the proportion of cells *(d)* S, *(e)* G1 and *(f)* G2 phases through time. Error bars correspond to the standard deviation between the 2 repetitions

To evaluate more precisely whether Tin2DN-induced cells simply proliferated more slowly or underwent sequences of growth arrest and regrowth we evaluated the amount of cells undergoing S phase during selected time periods using EdU pulses *(Figure 1b, c)*. In the non-induced group, the proportion of cells in S phase was stable, as the population grew exponentially. In the p16 induced cells, we observed a stable DNA synthesis arrest that was established within 48 hours. However in Tin2DN-induced cells, we observed 3 distinct phases that began with a slight decrease in the amount of cells that replicated their DNA, showing a primary proliferation reduction that coincided with primary telomere uncapping (Fig1S a,b). Then, after 72 hours of Tin2DN induction, we observed an increase in the number of cells undergoing S phase, indicating that the cells passed through the G1/S checkpoint despite harboring telomeric DDF (*Figure 2a*). Finally, the proportion of proliferating cells decreased steadily over the next few days and remained stable as cells entered senescence, as previously described for Tin2DN induction in normal cells ^25,26^.

**Figure 2:**
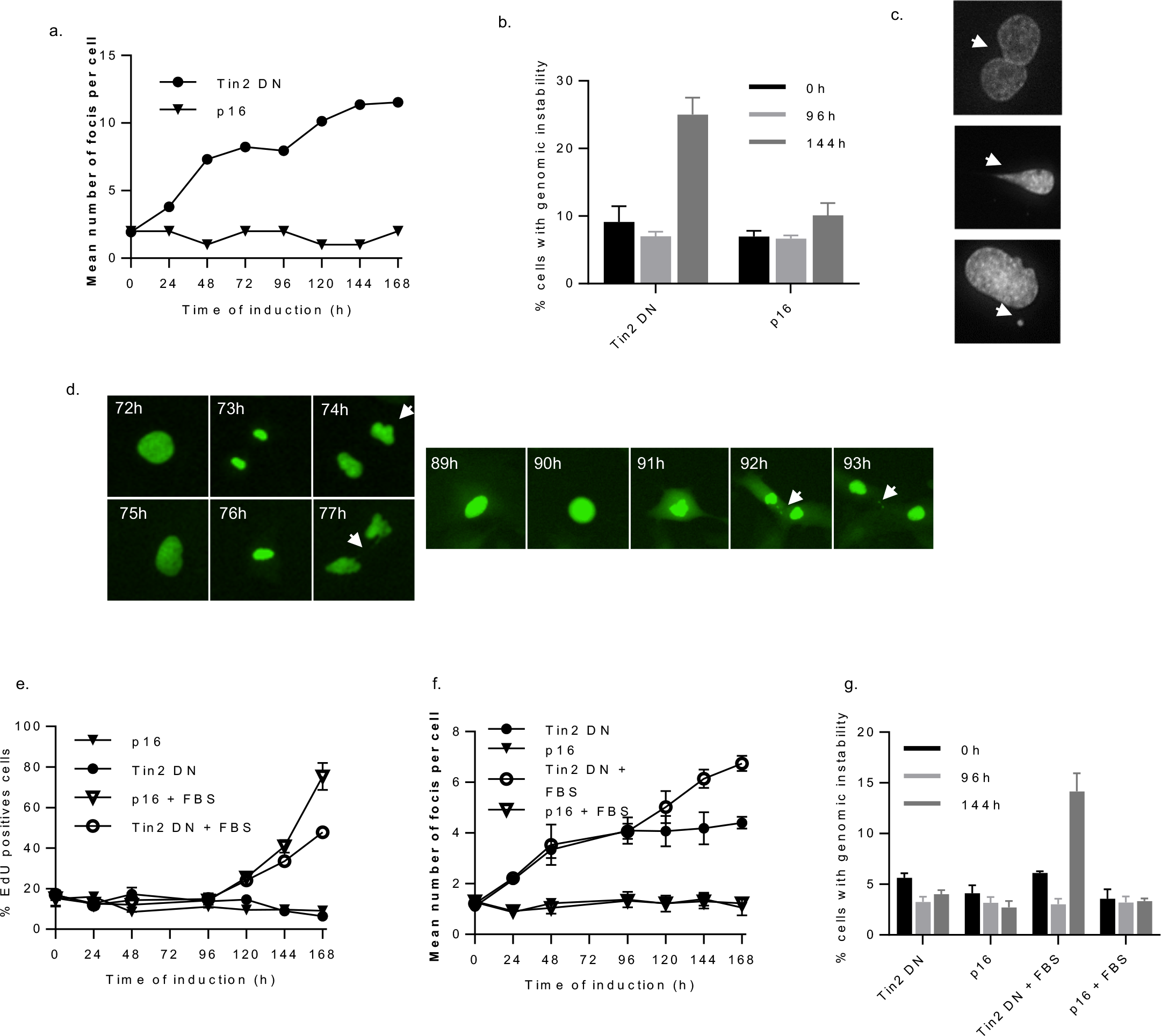
Cell division in the presence of dysfunctional telomeres causes secondary breaks and genomic instability. *(a)* Cells were treated with Doxycycline and fixed at different time points to perform immunofluorescence of 53BP1. Quantification of the mean number of foci per nuclei through time is shown. Error bars correspond to the standard deviation between the 5 repetitions *(b)* Quantification of the percentage of p16 or Tin2DN-induced cells showing at least one sign of genomic instability at different time points. We manually counted the number of cells showing either nuclei fusion, chromosome bridge, micronuclei or abnormal shaped nuclei. Error bars correspond to the standard deviation between the 5 repetitions *(c)* Examples of genomic instabilities observed after 144h of Tin2DN induction are shown with DAPI staining: nuclear fusion (top), chromosome bridge (middle) and micronuclei (bottom). *(d)* Pictures from live imaging of individual cells where Tin2DN was induced. The cells were pre-infected with H2B GFP to visualise DNA. Arrows are showing abnormal shaped nuclei (74h) chromosome bridge (77h) and micronuclei (92h-93h). *(e)* Cells were induced in serum starvation media (0.1%FBS) to prevent cell division. Cells re-proliferate when FBS is added to the media at 96h (+FBS). Error bars correspond to the standard deviation between the 3 repetitions *(f)* Quantification of the number of 53BP1 foci during low serum induction (0.1%FBS), or when serum is added to the media at 96h (+FBS) Error bars correspond to the standard deviation between the 3 repetitions *(g)* Quantification of the percentage of p16 or Tin2DN-induced cells showing at least one sign of genomic instability at different time points. We manually counted the number of cells showing either nuclei fusion, chromosome bridge, micronuclei or abnormal shaped nuclei during low serum induction (0.1%FBS), or when serum is added to the media at 96h (+FBS). Error bars correspond to the standard deviation between the 3 repetitions.

To further investigate cell cycle checkpoints bypassed in Tin2DN-induced cells we performed cell cycle FACS analysis during induction *(Figure 1d,e,f)*, which revealed that p16-induced cells accumulated in G1 in less than 48 hours, as expected for the action of this cyclin-dependent kinase inhibitor ^28^. Again, in Tin2DN-induced cells, we observed a complex multiple phase growth reduction process. The first phase involved a primary arrest with a decrease in the amount of S phase cells, and an increase in the amount of G1 and G2 cells. The second phase occurred after 72 hours of induction, where cells escaped G1 to enter S phase for approximately 48 hours. The final phase was an accumulation of the Tin2DN-induced cells in G1, with some cells remaining in G2. This last phase is consistent with the cell cycle distribution of primary HCA2 and BJ cells reaching replicative senescence ^29^, supporting entry into senescence.

### Bypass of the G2/M checkpoint in the presence of dysfunctional telomeres causes DNA breaks and genomic instability

We next quantitatively evaluated the amount of DSB-type DNA damage by quantifying 53BP1 DDF using immunofluorescence *(Figure 2a)*. Again, we observed 3 distinct phases throughout induction with Tin2DN. During the first 48 hours, cells showed an increase of DDF corresponding to the expression of Tin2DN and the opening of the t-loop, revealing the end of the chromosome. The number of DDF reached a plateau until the time point at 96 hours, followed by a second increase in DDF with a final plateau, presumably as a consequence of secondary breaks due to coincident G2/M checkpoint bypass. As the final increase in DDF coincided with growth recovery in these cells, we hypothesized that this increase was caused by the division of the cells with open telomeres, which could cause a high amount of genomic instabilities (GI) ^30^. We thus counted micronuclei, chromosome bridges and nuclear fusions *(Figure 2b and c),* and observed an increase in the percentage of cells presenting GI after 144 hours of Tin2DN expression/induction. Alternatively, no GI is observed in p16 induced-senescent cells nor after the primary Tin2DN-induced growth arrest *(Figure 2b)*. In order to confirm cell division-associated GI in real time, we then stably labelled the chromatin using H2B-GFP, which allowed us to follow cells in real time. We detected cells entering mitosis after 72 hours of induction, resulting in aberrant division as indicated by lagging chromosome, micro nuclei or mitosis duration above normal (figure 2d).

To confirm that the secondary breaks and GI were caused by cell division following checkpoint bypass, we prevented proliferation in Tin2DN induced cells using serum starvation (Figure 2e). Indeed, we did not observe any increase in the number of 53BP1 foci per cell nor GI when induced cells were kept in low serum *(Figures 2e and f)*. To clearly validate whether increased DDF and GI are triggered by cells division, we added serum back to starved cells, which immediately restored proliferation (*Figure 2e, + FBS)* leading to the previously observed phenotype of increased DDF and GI *(Figures 2f, g + FBS)*.

### Tin2DN induction results in a weak, differential DDR

Telomeric dysfunction has been shown to trigger growth arrest and senescence via the activation of an ATM-dependent ^31^ DDR signalling defined by p53 activation ^11^ and weak Chk2 activity ^31^. To validate this model, we probed the activity of these specific DDR events during primary telomeric dysfunction-induced growth arrest and secondary break-induced senescence. We also compared it to 1 Gy of X-ray irradiation (IR) that corresponds to 35 randomly distributed DSBs ^32^. This dose of irradiation is typically non-lethal and allows the cells to repair breaks and return to proliferation ^33^ as we observed in our system. We can therefore compare the strength of DDR activation following DNA DSB or telomere uncapping.

We observed two strong activations of ATM *(Figure 3a, arrow points at the bottom line),* after induction (time point 72h) and after the bypass (time point 168h) reflecting the 2 waves of damages: the primary telomeric uncapping that induced DNA damages, and secondary mitosis-induced DSB. The level of ATM phosphorylation was similar to the level obtained following 1Gy IR. However, we did not observe any significant activating phosphorylation of Chk2 following the primary response, whereas we detected strong phosphorylation following the bypass, at a similar level obtained by IR *(Figure 3a, c)*. Interestingly, results obtained with Chk2 could explain previously conflicting results in the literature, where previous observations showed no activation of Chk2 following telomere uncapping ^31^, while others showed that Chk2 was at least partially involved in the activation of replicative senescence ^34^. As the DDR cascade progressed, we observed that the low activation of pChk2 following Tin2DN induction (*Figure 3a, c*) leads to weak stabilisation of p53 (*Figure 3a, d*) and consequently, weak production of p21(*Figure 3a, e*). In contrast, the secondary breaks (occurring at approximately 120 h after induction, *Figure 2a*) resulted in strong activation of both Chk2, (*Figure 3a, c*) p53 (*Figure 3a, d*) and p21 (*Figure 3a, e*), at an amount similar to IR (*Figure 3a*). Therefore, two distinct waves of activation of the DDR cascade are demonstrated. The first one is exclusively caused by telomere uncapping, which is weak and does not include pChk2. The second activation is stronger and includes the phosphorylation of Chk2, triggered by secondary random breaks or GI.

**Figure 3:**
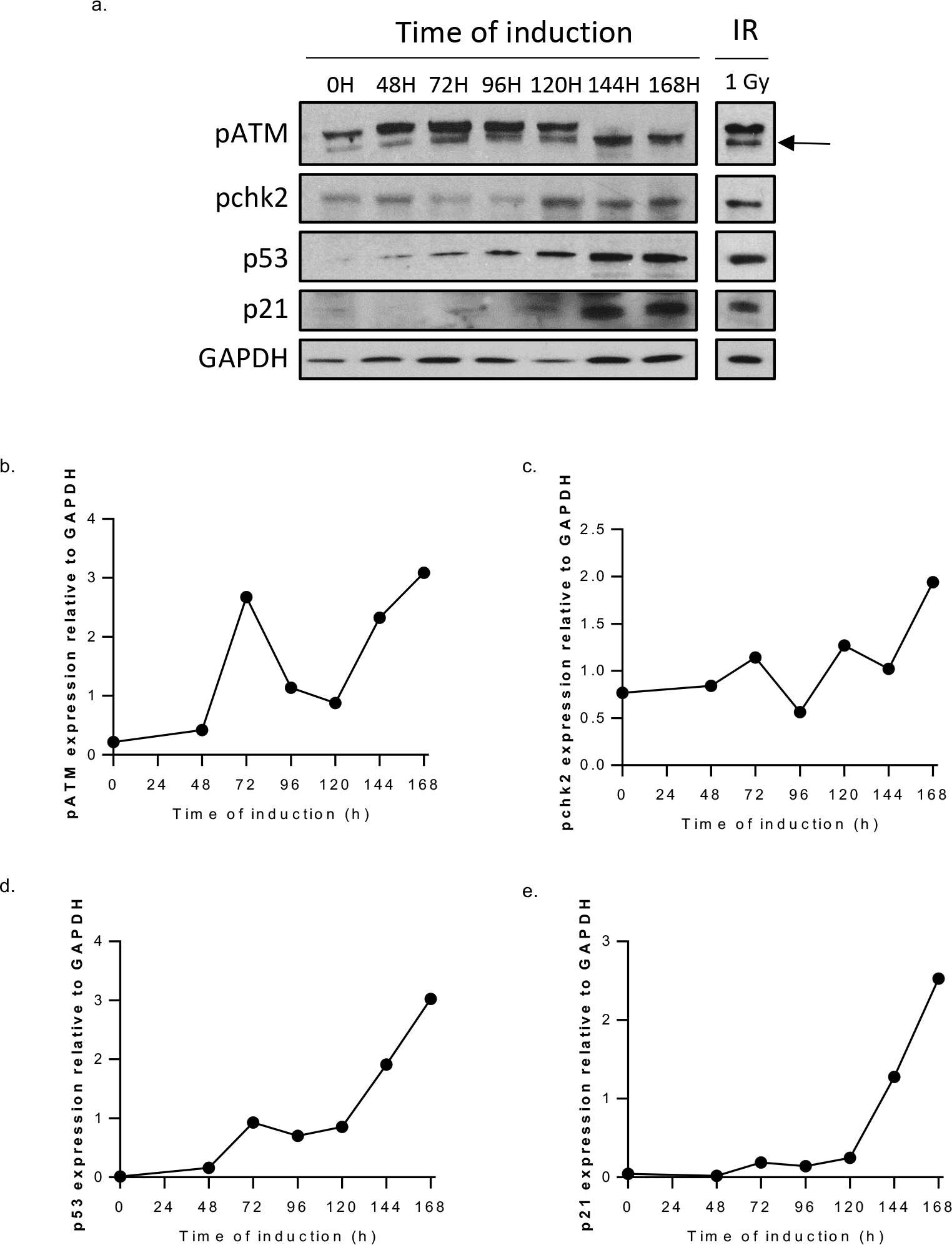
Telomeric dysfunction induces a weak DDR. *(a)* Cells were induced with Tin2DN and protein extracted at different time points. Western blot of `phosphorylated (p)?/activated?` DDR components throughout induction and at 4 h after 1 Gy X-Ray irradiation. Quantification of signal was performed with ImageJ and normalized to GAPDH levels for *(b)* pATM, *(c)* pChk2, *(d)* p53 and *(e)* p21. N=1

### Tin2DN-induced growth arrest is p53-dependent

Telomere uncapping has been shown to establish and maintain senescence via a DDR-p53-pathway ^11,12^. Despite the weak early p53 activation that we observed by western blot immediately following Tin2DN induction (*Figure 3a*), we directly tested the importance of p53 during multistep entry into telomere-mediated senescence. We infected our cells with a control shGFP or an shp53 lentivirus *(Figure 4a)* and performed another induction. We observed that p53 negative cells did not stop proliferating after the induction of Tin2DN *(Figure 4b),* and accumulated DNA damage *(Figure 4c)* and GI *(Figure 4d)* more rapidly, suggesting that p53 regulates the primary telomere-induced growth arrest. Accordingly, time-lapse experiments showed a very high frequency of mitotic abnormalities from the very beginning of the induction. Cells continued dividing, even though they were accumulating signs of damage. Mitoses were extremely long (over 3 hours), led to mitotic catastrophe and resulted in nuclei with lagging chromosome and of abnormal shape (*Figure 4e*). Eventually, after multiple days, most cells underwent mitotic catastrophe and detached from the dish. Similar results were obtain in previous studies, where p53 inactivated cancer cell lines were particularly sensitive to shelterin-induced telomere dysfunction and underwent rapid cell death, possibly via mitotic catastrophe caused by telomere dysfunction ^26^.

**Figure 4:**
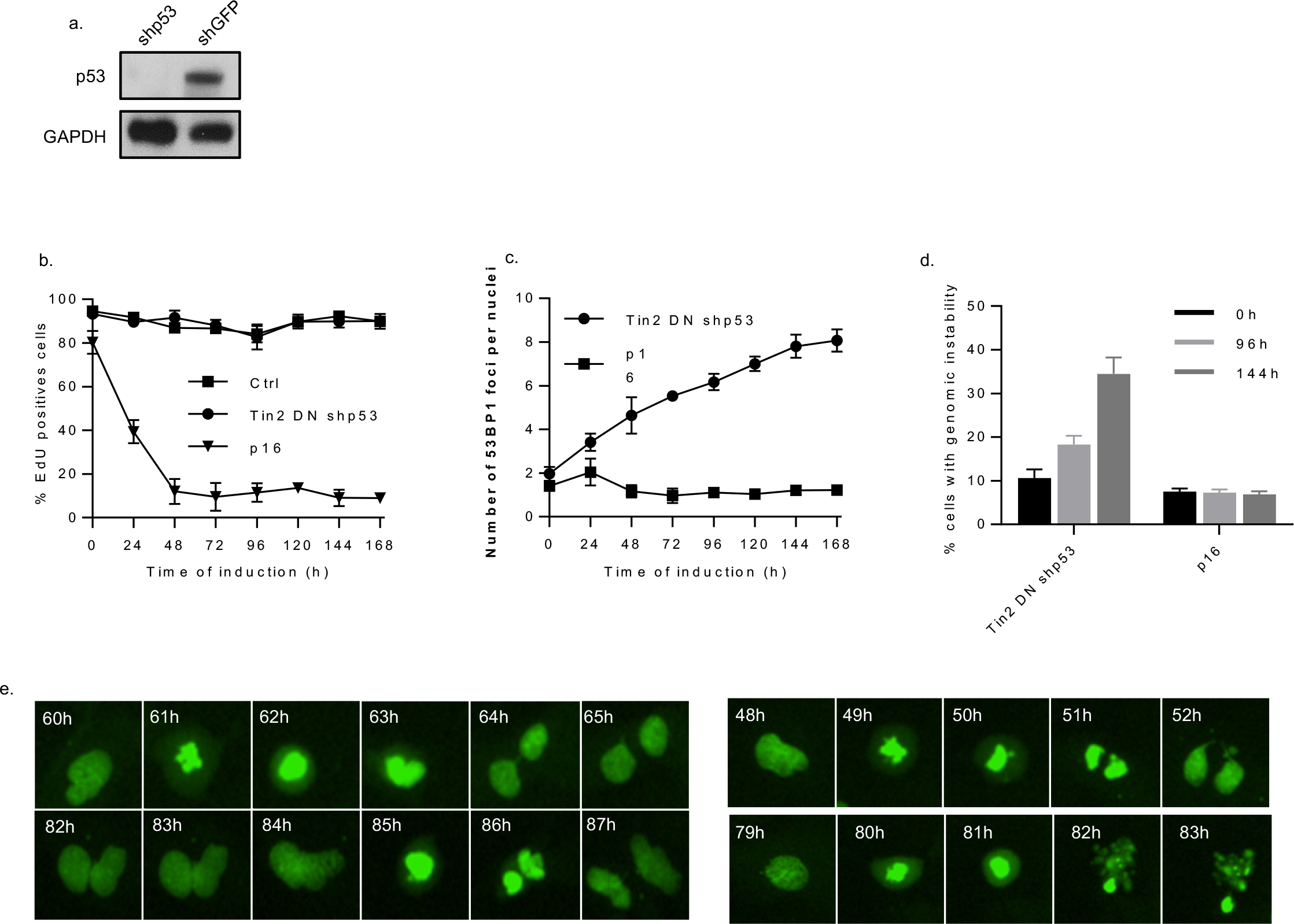
Tin2DN-induced growth arrest is p53-dependant. *(a)* BJ-U TR4s Tin2DNs cells were infected with an shp53 and, after selection, protein were extracted to validate the reduction of the level of p53 expression by western blot. *(b)* Cells were treated with Doxycycline and EdU was added to the media 24h prior to fixation. Cells were then stained and EdU positive cells quantified. Error bars correspond to the standard deviation between the 3 repetitions. *(c)* Cells were treated with Doxycycline and fixed at different time points to perform immunofluorescence of 53BP1. Quantification of the mean number of foci per nuclei through time is shown. Error bars correspond to the standard deviation between the 3 repetitions. *(d)* Quantification of the percentage of cells showing at least one sign of genomic instability. We manually counted the number of cells showing either nuclei fusion, chromosome bridge, micronuclei or abnormal shaped nuclei. Error bars correspond to the standard deviation between the 3 repetitions. *(e)* Pictures from live imaging of individual BJ-U TR4s Tin2DNs shp53 during induction. The cells were pre-infected with H2B GFP to visualise DNA. We can observe several examples of extended mitosis (3-4h long) resulting in chromosome bridges (arrows at 52h and 64h) micronuclei (arrow at 87h) or even mitotic catastrophe (82-83h).

### Tin2 depletion generates multistep entry into senescence

To ensure that our observations were not the result of a peculiar effect caused by the dominant negative mutation in Tin2DN, we reproduced our induction using shTin2. Inducible Tin2 depletion similarly triggered telomere dysfunction that induced a primary arrest (*Figure 5a*), and secondary mitosis-induced DNA breaks (*Figure 5b*) that gave rise to GI and a stable senescence growth arrest (*Figure 5c*).

**Figure 5:**
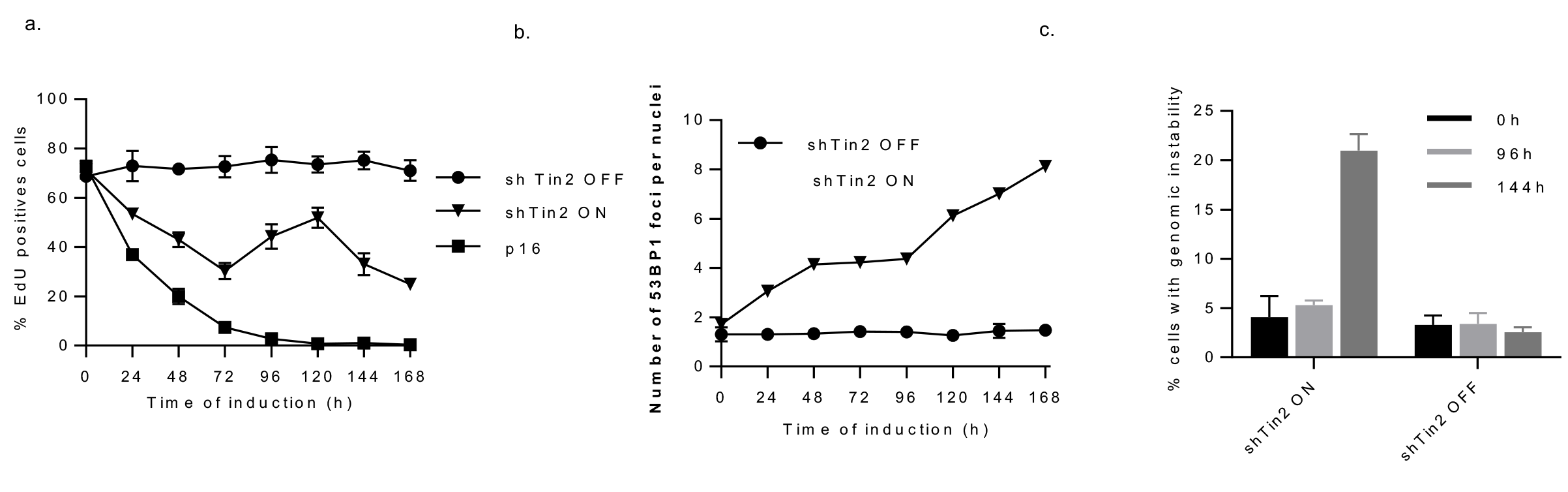
Results are reproducible with shTin2. *(a)* Cells were treated with Doxycycline and EdU was added to the media 24h prior to fixation. Cells were then stained and EdU positive cells quantified. Error bars correspond to the standard deviation between the 3 repetitions. (b) Cells were treated with Doxycycline and fixed at different time points to perform immunofluorescence of 53BP1. Quantification of the mean number of foci per nuclei through time is shown. Error bars correspond to the standard deviation between the 3 repetitions. *(c)* Quantification of the percentage of p16 or Tin2DN-induced cells showing at least one sign of genomic instability at different time points. We manually counted the number of cells showing either nuclei fusion, chromosome bridge, micronuclei or abnormal shaped nuclei. Error bars correspond to the standard deviation between the 3 repetitions.

## DISCUSSION

### A new model for replicative senescence induction

By inducing telomeric dysfunction under controlled and specific conditions, we showed that dysfunctional telomeres are not direct inducers of senescence in normal human cells. Previous knowledge concerning entry into telomere-mediated replicative senescence stipulates that replicative senescence is caused by the direct activation of the DDR following the uncapping of telomeres. Alternatively, our model system showed an indirect mechanism where dysfunctional telomeres trigger senescence via the induction of secondary DNA breaks and GI. We find that the expression of Tin2DN uncapped telomeres and created a DDR with the formation of 53BP1 DDF as expected *(Figure 1S)*. However, this telomere dysfunction-induced DDR was slightly different ^31^ and not as strong when compared to a DSB-induced DDR initiated by radiation. Indeed, we observed no activation of pChk2, and a weak activation of p53 and p21 *(Figure 3)*. We speculate that these differences are the cause of the instability of the Tin2DN-induced growth arrest *(Figure 1)*. The attenuated activation of cell cycle checkpoints following Tin2DN induction resulted in cells re-entering the cell cycle and dividing, despite the presence of dysfunctional telomeres. It has been well described that under many conditions a cell dividing with damaged DNA is likely to suffer from abnormal mitosis and GI ^35,36^. This is what we observed in our system, with an increase in the number of 53BP1 DDF, and the appearance of signs indicating GI. We also confirmed that these abnormalities were related to mitosis as blocking cell division was sufficient to prevent GI or any increase in the number of DDF. These results showed that telomere uncapping leads to an unstable growth arrest that will cause secondary breaks and GI, which will lead to senescence (*Figure 6*). It is interesting to note that this slight increase alone (from 4 to 8 foci per cell) is enough to establish a stable growth arrest, supporting the idea of a threshold of DDR. We could also speculate that the cells with the highest amount of damage died of mitotic catastrophe. Importantly, a high frequency of polyploidy and multiple chromosomal abnormities have been reported in cells approaching senescence in several distinct studies ^37–40^. This suggests that our observations are not confined to our system, but that our proposed multistep telomere-mediated senescence model may occur in natural aging through telomere shortening.

**Figure 6:**
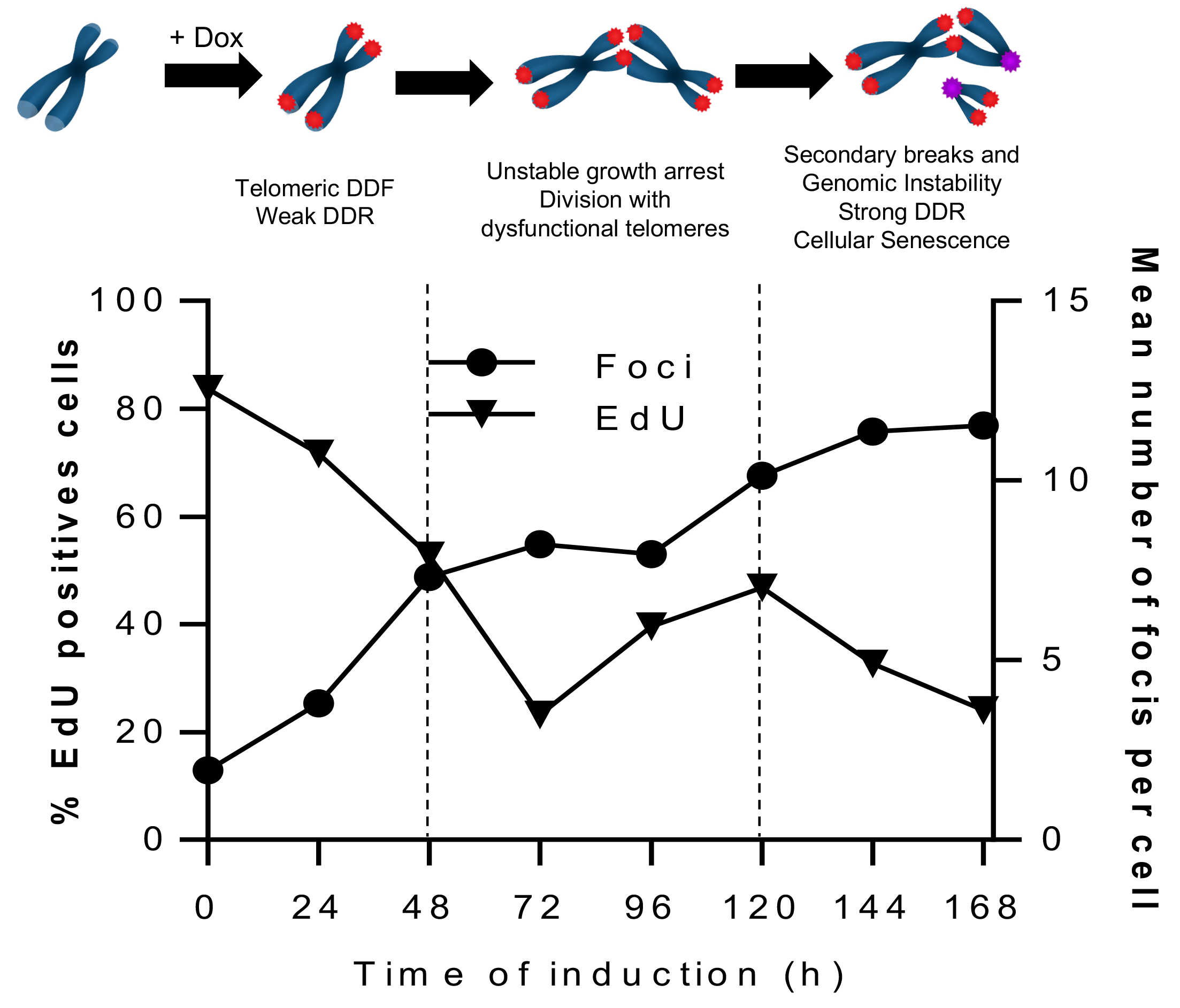
A new model for entry into senescence following telomeric dysfunction. This model is based upon our results which show alignment between the time-course of EdU proliferation and the timeline of 53BP1 DDF formation. We conclude that the expression of Tin2DN induced by Doxycycline treatment (+ Dox), leads to a weak DDR with telomeric DDF formation but unstable growth arrest. As the telomeres are uncapped, there is a high susceptibility for inter-chromosomal fusions. Although we did not observe them directly, the high amount of GI following divisions is evidence of their occurrence. Cell division with dysfunctional and potentially fused telomeres leads to randomly distributed, secondary DSB that will trigger a strong DDR and a stable senescence associated growth arrest.

We hypothesize that the lack of stability may come from the limited activation of p53 and p21, however, when we repeated our time course induction in shp53-infected Tin2DN-induced cells, we observed no proliferation arrest with an accumulation of GI and a constant increase in the number of 53BP1 DDF. This showed that p53 activation, as weak as it may have been, was absolutely essential for primary growth arrest.

Altogether, these results propose a new model for replicative senescence, where the permanent growth arrest is not directly caused by the uncapped telomeres, but by the secondary breaks caused by the cell cycle checkpoints adaptation.

### Loss of TRF2 might causes chromosome fusions

This conclusion proposed by our model may be unexpected and counter-intuitive since allowing cell division in the presence of DNA damage is a risk for inducing cancer. The evolutionary gain of this model is therefore questionable. Is DNA damage tolerance a flaw or an active way of ensuring a permanent growth arrest? How does the cell ensure the balance between survival and genome stability?

A hypothesis would be that the loss of Tin2 function via the dominant negative mutation results in the loss of TFR2 as suggested by Kim et al. ^25^ and creates uncapped fusogenic telomeres ^4,41^. The inhibition of DDR would therefore be lost, allowing fusions between chromosomes that could be interpreted by the cell as a DNA repair. The cell would then deactivate its checkpoints, re-enter into the cell cycle with fused telomeres, generating new damage. Nonetheless, we do not have any evidence that a similar consequence is taking place when we induce the expression of shTin2, although we observed exactly the same phenotype in both cases.

This model would be another verification of the importance of DDR inhibition at the telomeres, as it is the repair itself that causes new damage and GI. It is possible that this tolerance is an active mechanism, created to amplify the damages that would promote a strong proliferation arrest.

### An example of checkpoint adaptation in mammals?

In yeast, cell cycle checkpoint adaptation allows the cell to re-enter into cell cycle and division in the presence of DNA damage in order to split the DNA lesions between the resulting two daughter cells, via an asymmetric division ^42^. This guarantees the survival of at least one cell, while the other daughter cell dies of mitotic catastrophe. In multicellular organisms, the DNA damage adaptation could be a way to bring the cell into a cell phase that is more appropriate for repair ^43^, or a preliminary step that leads to cell death by mitotic catastrophe ^44^. In this context, the cell cycle checkpoint bypass in the presence of DNA breaks would be a way of amplifying the damages, ensuring a subsequently strong proliferation arrest.

Another example is observed during DNA replication ^45^ where the DNA damage tolerance pathway or Translesion synthesis (TLS) is a process that allows DNA synthesis in the presence of DNA breaks. This error-prone mechanism prevents the block of the replication fork and allows the cell to continue through the cell cycle despite damage ^46^.

Although, the maintenance of genome integrity is crucial to prevent mutations and cancer development, a cell cannot constantly stop the cell cycle at any sign of stress. The existence of several tolerance pathways allows the cell to continue its division in case of minor damage ^46^. In the case of telomeres, it is probably counteractive for cells to have telomeres detected as DNA damage, as it may introduce inappropriate fusions and recombination, specially during replication when the opening of the t-loop is inevitable. Hence, the t-loop serves to shield and protect the chromosome end, and the active inhibition of DNA damage detection and repair pathways by the proteins of the shelterin complex, prevents an excessive response from the cell. However, these tolerance mechanisms/strategies present a risk in that any dysfunction in another DDR pathway, such as p53, could lead to a very high amount of GI and could eventually create mutations that develop into cancer ^30,47,48^.

In this study, we provide the first direct evidence of telomeric dysfunction tolerance in normal human cells. As cells divide with uncapped telomeres, they create new DSB that eventually lead to a permanent senescence associated cell growth arrest. We propose a new model for entry into replicative senescence, offering a new explanation for aging-associated genomic damages and instabilities. Our results could also provide clues to explain how a cell can escape cellular senescence and transform into cells that become cancerous.

## ACKNOWLEDGEMENTS

We thank members of the Rodier laboratory for valuable comments and discussions. We thank Jacqueline Chung for valuable comments and English editing. This work was supported by the Institut du cancer de Montréal and by grants from the Canadian Institute for Health Research [MOP114962]. FR is supported by the Fonds de Recherche Québec Santé junior I career award [22624]. SG is supported by an Institut du cancer de Montréal Canderel scholarship.

---

**Figure S1:**
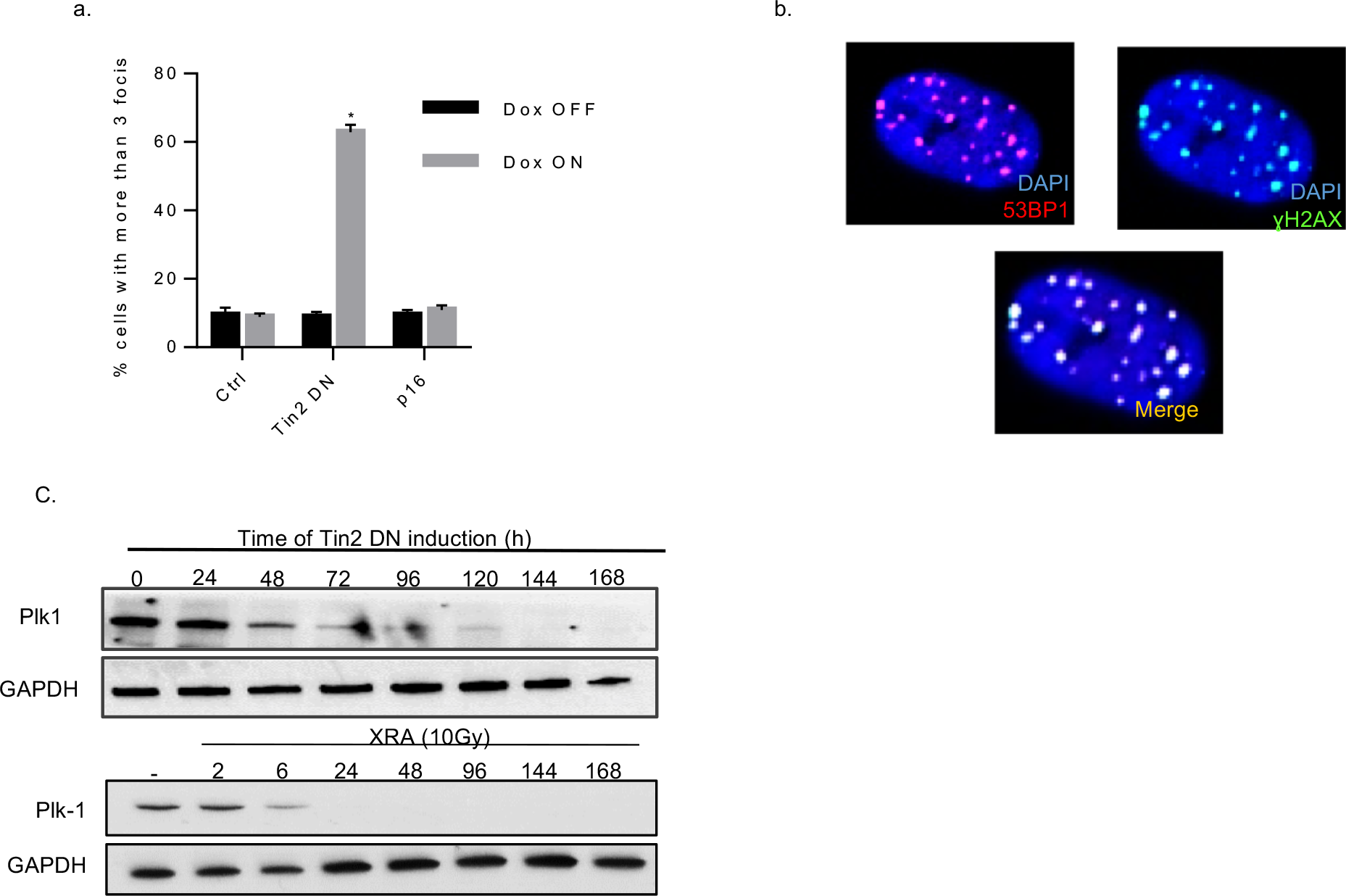
Tin2DN expression induces DNA damage foci formation. *(a)* Cells were treated, or not (Ctrl), with Doxycycline and fixed after 48h to perform immunofluorescence of 53BP1. Quantification of the mean number of foci per nuclei is shown. *(b)* Representative picture of BJ-U TR4s Tin2DNs cells induced for 48h and immunostainned for 53BP1 (red) and γH2AX (green). Although we did not perform it ourselves, a previous study using the same truncated form of Tin2DN showed that the DNA damage foci were localised at the telomeres (Kim et al., 2004, 2008).

